# Swimming ability of the *Carybdea marsupialis* (Cnidaria: Cubozoa: Carybdeidae): implications for its spatial distribution

**DOI:** 10.1101/2023.05.06.539705

**Authors:** Cesar Bordehore, Sara Manchado-Pérez, Eva S. Fonfría

**Affiliations:** Multidisciplinary Institute for Environmental Studies (IMEM) “Ramon Margalef”, University of Alicante, Campus San Vicente del Raspeig s/n, 03690 San Vicente del Raspeig, Spain; Department of Ecology, University of Alicante, Campus San Vicente del Raspeig s/n, 03690 San Vicente del Raspeig, Spain; Department of Ecology and Hydrology, University of Murcia, Campus Espinardo s/n, 30100 Murcia, Spain

**Keywords:** *Carybdea marsupialis*, Advection, box jellyfish, Effective Displacement Index, surface currents, swimming speed

## Abstract

Although usually considered part of the plankton, cubozoans are strong swimmers. The aim of this study was to determine the influence of the active swimming ability of the box jellyfish *Carybdea marsupialis* on the spatial distribution of a well-studied population off Dénia (NW Mediterranean) where adults and juveniles do not overlap geographically. To achieve this aim, we analyzed the swimming speed, proficiency, effective velocity, and effective displacement index (EDI) of 27 individuals with diagonal bell widths (DBW) ranging from 1.1 to 36 mm. The laboratory analysis utilized conventional video recordings and the video analysis tool Tracker. Mean swimming speed for small juveniles (DBW ≤ 5 mm), medium juveniles (5 mm < DBW< 15 mm) and adults (DBW ≥ 15 mm) were 9.65 ± 0.76 mm^-1^, 21.91 ± 2.29 mm^-1^ and 43.10 ± 1.78 mm^-1^ (mean ± s.e.m.), respectively. Comparing these results with the local currents obtained from drifting buoys analyzed in the area over the course of three years, adults would be able to swim strongly enough to overcome almost 70% of the currents, whereas the small juveniles would not reach 17%. This allows larger individuals to select their habitat, while smaller individuals are left dependent on advection. Although experiments adding currents in aquaria would be necessary to confirm these theoretical results, the data obtained would be useful in improving the performance of bio-mathematical models used to predict jellyfish blooms since, even though the sting of *C. marsupialis* is non-fatal, it may produce systemic effects in sensitive swimmers.

**Summary statement:** The analysis of the swimming ability of *C. marsupialis* elucidates its key role in the spatial distribution of a northwestern Mediterranean population.

## INTRODUCTION

The word “plankton” is a collective term for pelagic organisms adapted to life in suspension, which have in common the passive entrainment by water currents (Reynolds and Padisák, 2013). Historically, jellyfish of the classes Scyphozoa and Cubozoa have been classified in this category, but some studies indicate otherwise. Studies have demonstrated that the scyphozoans *Aurelia* sp., *Cyanea capillata, Phacellophora camtschatica, Rhizostoma octopus, Rhizostoma pulmo* and *Rhopilema nomadica* can actively swim against a current and exhibit positive rheotaxis (Fossette et al., 2015; Malul et al., 2019; Moriarty et al., 2012; Rakow and Graham, 2006). In addition, other movements such as vertical migrations or sun-compass horizontal migrations have also been reported (Albert, 2007; Hamner and Hauri, 1981; Hamner et al., 1994; Hays et al., 2012). Cubozoans even go a step further. In addition to swim at counter-current (described for *Tripedalia cystophora* and *Chiropsella bronzie* (Garm et al., 2007), *Chironex fleckeri* (Schlaefer et al., 2018) and *Copula sivickisi* (Schlaefer et al., 2020)), their unique visual system, which comprises a total of 24 eyes (Garm and Bielecki, 2008), allows them to direct their swimming in response to external stimuli (Kingsford et al., 2021). Obstacle avoidance has been reported for *C. fleckeri* (Hamner et al., 1995; Schlaefer et al., 2018) and *T. cystophora* and *C. bronzie* (Garm et al., 2007). Positive phototaxis, the tendency to move towards a light source, has also been observed in *T. cystophora* (Buskey, 2003) and *C. sivickisi* (Garm et al., 2012), which is a common trait among cubozoans and is often used to collect them during nocturnal samplings (Acevedo, 2016; Bolte et al., 2021; Morandini et al., 2014). All these findings suggest that, at least for adult specimens, it may be more appropriate to classify them as part of the nekton, a term that includes pelagic organisms that swim freely independently of currents.

In the Mediterranean Sea, *Carybdeda marsupialis* (Linnaeus, 1758) is the sole reported species of the Class Cubozoa (Acevedo et al., 2019). Usually observed in low densities, over recent decades it has been massively detected in some Italian (mainly in the Adriatic Sea), Tunisian and Spanish coastal zones (Boero, 2013; Bordehore et al., 2011; Gueroun et al., 2015). Likeother cubozoan jellyfish, *C. marsupialis* has a biphasic life cycle with a benthic polyp and a free-swimming medusa. Recently detached medusae measure less than 2 mm DBW (Diagonal Bell Width, distance between opposite pedalia at level of pedalia joining bell) while adults can grow up to 40 mm (Acevedo et al., 2019). On the Spanish coast of Dénia, where it has been detected in higher abundance and where a population is maintained year after year, *C. marsupialis* shows a clear seasonality. Polyp metamorphosis occurs in spring (Acevedo et al., 2019), resulting in the appearance of small juveniles from May to July (DBW <5mm), medium-size juveniles (5 ≤ DBW <15mm) from July to August and adults (DBW >15mm, up to 40 mm) approximately from August to October or early November (Acevedo, 2016; Bordehore et al., 2015a; Bordehore et al., 2020a; Canepa et al., 2017). *C. marsupialis* shows a clear coastal distribution at all stages (e.g. 90.8 % of the total catches of the samplings carried out in the period 2013-2015 correspond to the strip located between 0 and 15 m from the coast (Bordehore et al., 2020b), however, adults tend to concentrate in areas with high food availability (high levels of primary and secondary production), while small-juveniles can be found in other locations (Bordehore et al., 2020a; Canepa et al., 2017). The limited dispersal of C. marsupialis may be attributed to coastal dynamics and/or its ability to actively maintain itself in a chosen area by swimming.

Thus, studying the swimming ability and current resistance across different developmental stages is crucial in understanding the species’ spatial distribution.

To obtain swimming behavioral data on jellyfish within their habitats, electronic tags (acoustic transmitters and acceleration data loggers) have been used (Fossette et al., 2015; Gordon and Seymour, 2009; Moriarty et al., 2012). Nevertheless, the use of these tags is limited by the size of the species, among other constraints (Fossette et al., 2016). In our case, adults of *C. marsupialis* reach a maximum of 40 mm in DBW (half the size recommended by Gordon and Seymour 2009 to reduce potential confounding effects due to tag weight), making it necessary to develop smaller transmitters, which would be impossible to place in the case of early (5 mm) or late (15 mm) juveniles.

The main objective of this study was to elucidate the influence of active swimming ability on the spatial distribution of *C. marsupialis*. The specific goals include: i) To determine the ontogenic swimming speed and the effective displacement of *C. marsupialis* in the laboratory using video recordings and free tracking software, and ii) To provide data on their theoretical ability to overcome currents.

## MATERIALS AND METHODS

### *Carybdea marsupialis* collection and care

Cubomedusae were collected between June and October 2016 on the coasts of Dénia and El Campello (Spain, Western Mediterranean Sea) by day (for juveniles and adults specimens) and night (only for adults) samplings (Fig. 1). Diurnal samplings were performed on foot, walking for a duration of 15 min parallel to the coastline within 15 m from the shoreline at 0.4 m/s using hand plankton nets (length 1.5 m, mouth area 0.15 m^2^, mesh size 500)(Bordehore et al., 2020a). At night, waterproof LED lights (50W, 12 V) were used. Two lights were mounted at dusk, 15 meters from the shoreline and 1 meter deep. After 1 hour of light exposure, the first cubozoans appeared and they were gathered using plastic beakers (Bordehore et al., 2020b). In both types of samplings, once collected, *C. marsupialis* individuals were placed into 50 L plastic containers filled with clean seawater and transported to the Dénia Montgó Scientific Station, where they were maintained in aquaria with filtered seawater at 24.5 ºC and a salinity of 37.4. They were fed daily with *Artemia salina* nauplii.

**Fig. 1.**
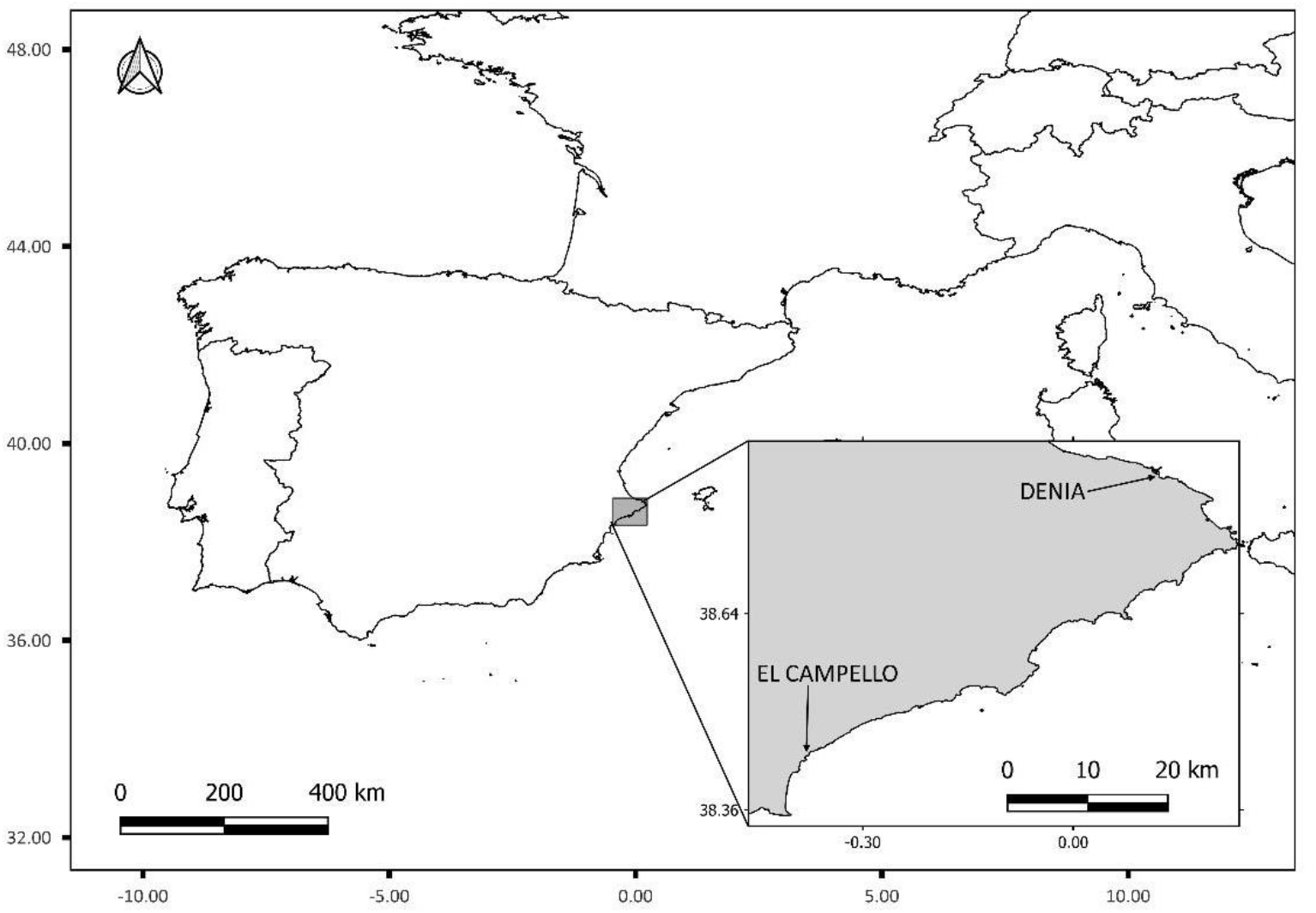
Sampling areas (Dénia and El Campello) on the Spanish coast (Western Mediterranean Sea).

In the laboratory, the DBW of each specimen was measured using a stereoscopic microscope (LEICA S8APO) for medusae that were ≤ 10 mm wide and with calipers for larger ones. Afterwards, cubozoans were grouped into three categories that represented different developmental stages based on their DBW: small (≤ 5 mm), medium (between 5 and 15 mm) and large (≥ 15 mm) individuals (Acevedo et al., 2013).

### Surface currents speed

Within the framework of the LIFE CUBOMED project (www.cubomed.eu), periodic samplings were carried out between 2013 and 2015 along the 17 km of the coast of Dénia at 0, 250, and 500 meters from the shoreline. Surface currents speeds were obtained by deploying two drift buoys at different sampling locations (11 points at 0 m and 4 points at 250 and 500 meters) and recording their initial and final waypoints (5 min after release) with a GPS device (Garmin 72H)(Bordehore et al., 2020a; Canepa et al., 2017). The speed of the buoys were calculated from geographic coordinates and using the Haversine mathematical formula and the deployed time.

### *Carybdea marsupialis* swimming experiments

A total of 27 specimens (12 small juveniles, 7 medium-size juveniles and 8 adults) were evaluated. The DBW ranges for the three classes were: 1.1 to 3.9 mm for small, 5.4 to 14.2 mm for juveniles and 29.8 to 36.0 mm for adults. Individuals were video recorded within 48 h of collection while freely swimming in different recipients significantly larger than their body size. A minimum aquarium volume to jellyfish volume ratio of >3000 was ensured for all classes to avoid wall effects or other artifacts of the laboratory environment (Dabiri et al., 2010).

The small juveniles were analyzed in a Petri dish of 8.6 cm in diameter (equivalent to a range of 1/78.2 – 1/22.1 DBW/petri dish) and 1.2 cm in height filled with 44 mL of filtered seawater at 50 μM. Lateral face was covered with black tape to avoid the incidence of lateral light and the possible positive phototropism effects that *C. marsupialis* exhibits even in small sizes (Bordehore, 2014). Medium-sized specimens were divided into two subgroups (DBW >10 mm and DBW<10mm) and studied in two different cylindrical aquaria, the larger one of 29.5 cm diameter x 19.1 cm height, with 10 L of filtered seawater, and the smaller one of 17.3 cm diameter and 19.9 cm height, with 1.2 L. The minimum ratio between the DBW of the specimens and the diameter of the container was 1/28.6 and 1/32.0, respectively. Adults (>1.5 cm DBW) were placed in a rectangular aquarium (26 cm width x 29.5 cm height x 58 cm length) with 25L of filtered seawater. In this case, the length was 19.5 times larger than the diameter of the smallest adult tested (DBW=29.8 mm), respectively.

The Petri dish and 1.2 L aquarium were recorded using a Hercules Dualpix Infinite Camara in zenith position, and fixed on a tripod. For the 10 L and 25 L aquariums a Sony Handycam DCR-SR32 was used in the same position. To ensure that perpendicular trajectories to the main camera were recorded, an auxiliary camera (GoPro Hero3+) on the side of the aquariums was utilized. In all cases, a grid was included as a reference in the background, including the Petri dish.

#### Video image processing

The GoPro Hero+3 images were analyzed using GoPro Studio software (goo.gl/DiruZY) to eliminate the fisheye effect that alters the trajectory.

In the adult recordings, sections where the jellyfish remained in the corners of the aquarium or where their movement was not horizontal (i.e. perpendicular to the main camera) were eliminated using editing software Movie Maker and Sony Vegas Pro 13 (goo.gl/xLpB9n).

#### Swimming kinematics

We analyzed swimming speed (distance/time), displacement and effective velocity (defined here as total displacement/time) for each individual using the free software video analysis and modeling tool Tracker (https://physlets.org/tracker/).

Considering speed as the time rate at which an object is moving along a path, swimming speeds of *C. marsupialis* were calculated from the position-time data of the reference point in the apex of the bell (Rubio-Tortosa et al., 2016). Tracker’s software uses the finite algorithm shown below (eqn 1 and 2), which defines the parameter evaluated for a step as the average value over 2-step intervals. Subscripts refer to step numbers, and dt is the time between steps in seconds.

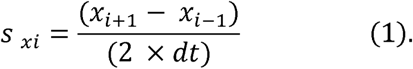

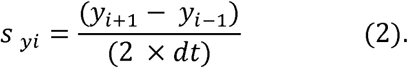

A minimum of 50 intervals were analyzed per specimen (with a maximum of 438 and a mean ± s.e.m., standard error of the mean, of 200 ± 20), with a 4-5 step size (number of frames per interval).

To determine displacement and effective velocity, 5 segments were selected randomly for each specimen. Displacement was measured as the linear difference between the final position and its starting position, using the Tracker’s tape measure. Knowing the time elapsed between the two points, effective velocity was calculated.

To represent the relationship between effective velocity and swimming speed, we created an index called “Effective Displacement Index” (EDI), obtained by dividing the velocity by the speed. It can range from 0 to 1. When EDI = 1, speed and velocity have the same value meaning that the jellyfish traveled in a straight line (distance and displacement are equal). Values of EDI close to 0 indicate that the jellyfish do not follow a linear path, but a more complex trajectory (in this case distance is greater than the displacement)(Fig. 2).

**Fig. 2.**
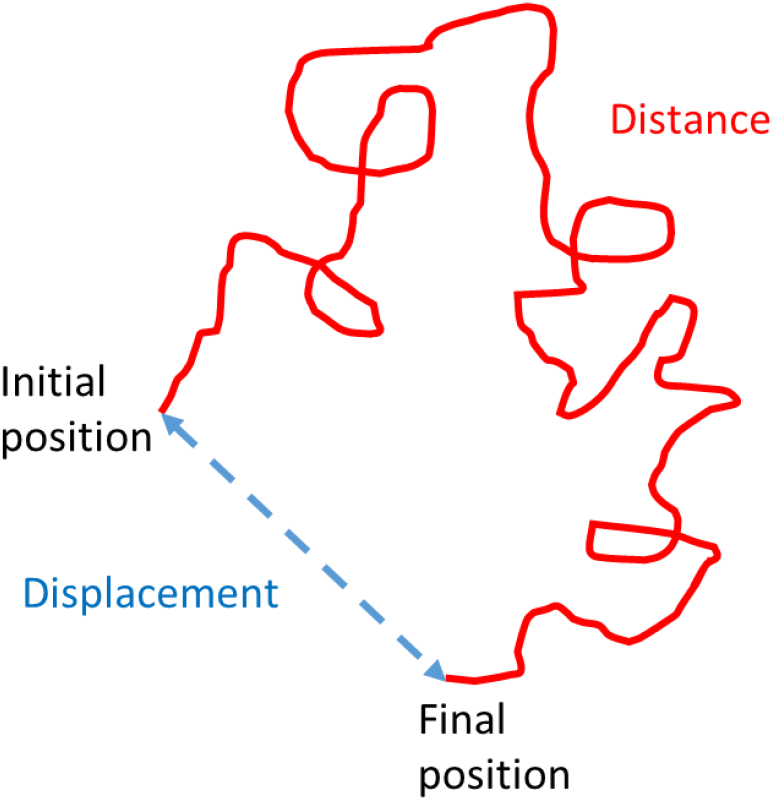
Difference between distance and displacement (and, consequently, for the same elapsed time, speed and velocity) of a moving object.

The proficiency of swimming was quantified as the diameter-normalized swimming speed, as follows (Eqn 3):

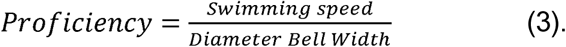

## RESULTS

### Swimming speeds and effective velocities

In the laboratory conditions specified, the average swimming speed of all *C. marsupialis* specimens (N=27) was 22.74 ± 2.89 mm s^-1^ (mean ± s.e.m.). It increased proportionally to DBW (Fig. 3). Per groups, the smaller ones (1.1 ≤ DBW ≤ 3.9 mm) swam at 9.65 ± 0.76 mm s^-1^ (mean ± s.e.m., n=12), whereas the medium ones (5.4 ≤ DBW ≤ 14.2 mm) did it at 21.91 ± 2.29 mm s^-1^ mm s^-1^ (mean ± s.e.m., n=7), and the adults (29.8 ≤ DBW ≤ 36 mm) at 43.10 ± 1.78 mm (mean ± s.e.m., n=8).

**Fig. 3.**
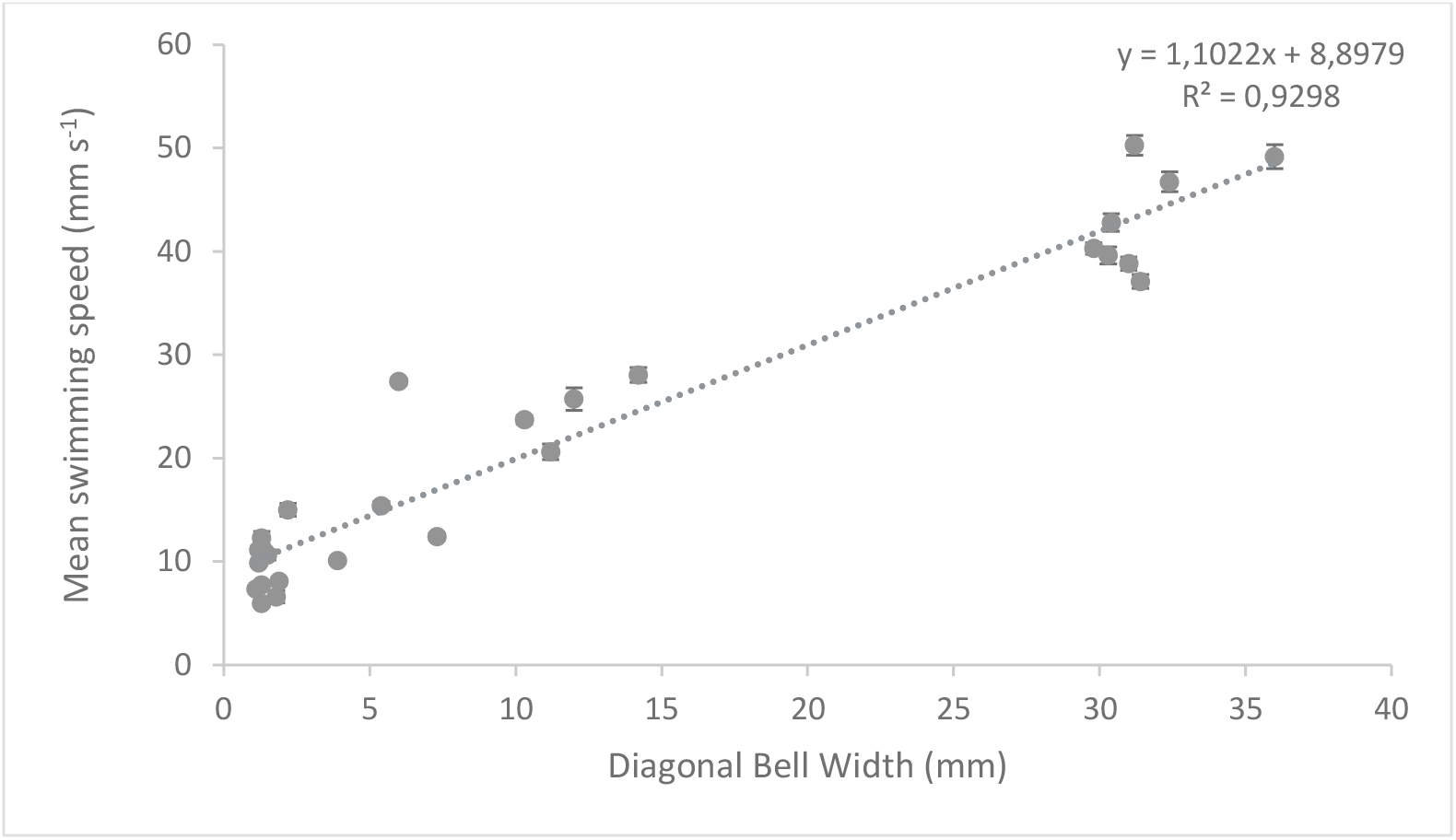
Mean swimming speed of *C. marsupialis* with increasing size (mean ± s.e.m.) (n=27).

Regarding the average maximum velocity, it also varieswith the size (Fig. 4), being 41.23 ± 4.93 mm s^-1^ (mean ± s.e.m.) when all the medusae were grouped together, and 18.64 ± 1.34 mm s^-1^, 40.98 ± 4.34 mm s^-1^ and 75.35 ± 3.17 mm s^-1^ when divided by size class, from smallest to largest, respectively (mean ± s.e.m., same N per group as previous analysis). The range of maximum and minimum individual velocities was as follows: 0.04-29.75 mm s^-1^ for smaller jellyfish, 0.67-54.27 mm s^-1^ for medium-size specimens and 5.72-87.23 mm s^-1^ for adults.

**Fig. 4.**
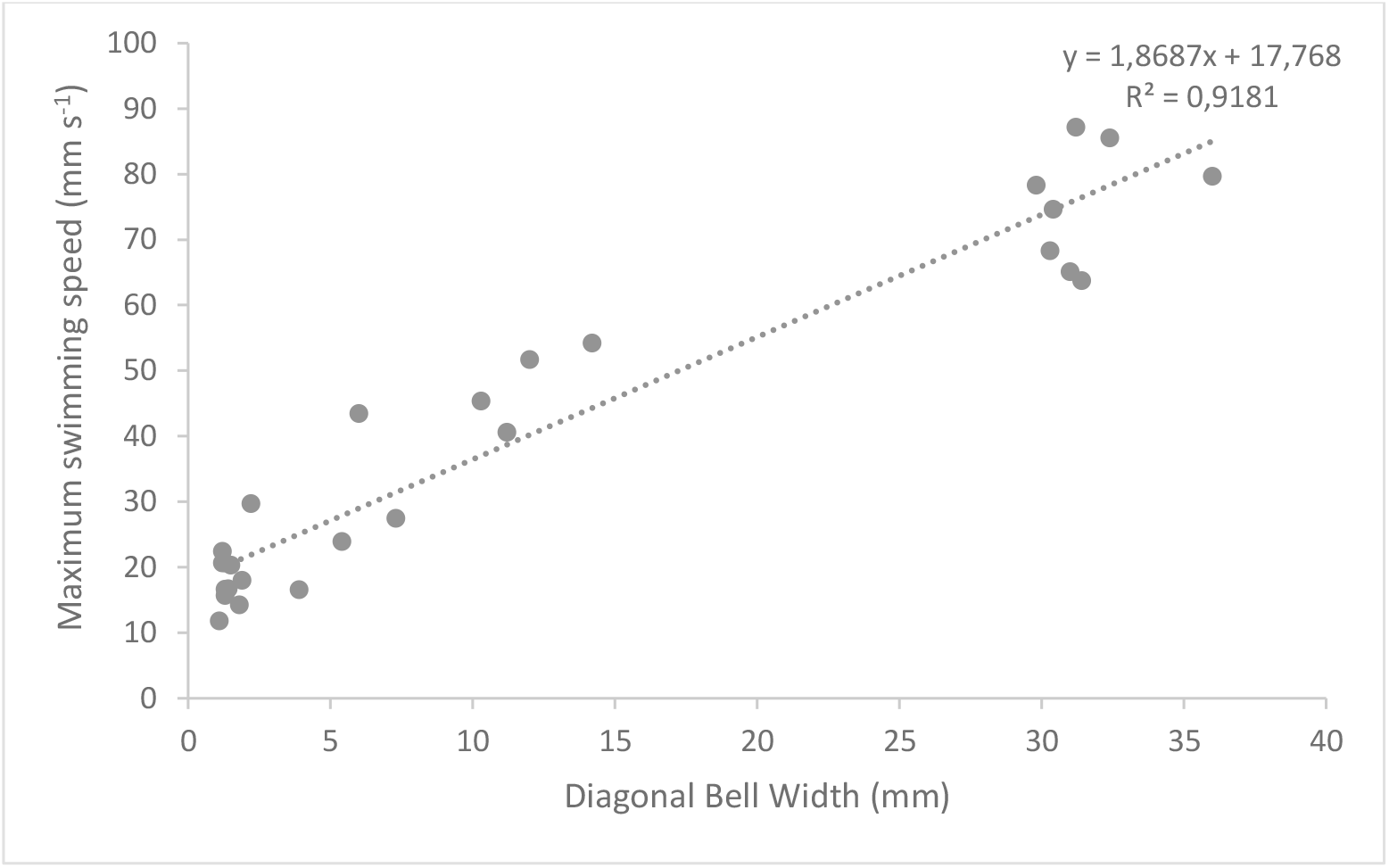
Maximum swimming speed of *C. marsupialis* with increasing size (n=27).

The effective velocity was also proportional to the DBW (Fig. 5). It was +5.04 ± 0.68 mm s^-1^ for juvenile 1, +18.83 ± 2.54 mm s^-1^ for juvenile 2 and +38.83 ± 3.13 mm s^-1^ for adults (mean ± s.e.m.). The calculated EDI for each group was 0.51 ± 0.0.5, 0.84 ± 0.06 and 0.90 ± 0.05 (mean ± s.e.m.), respectively.The trajectory of small juveniles tends to be spiral-shaped, while medium-size specimens and adults tend to have a straighter trajectory.

**Fig. 5.**
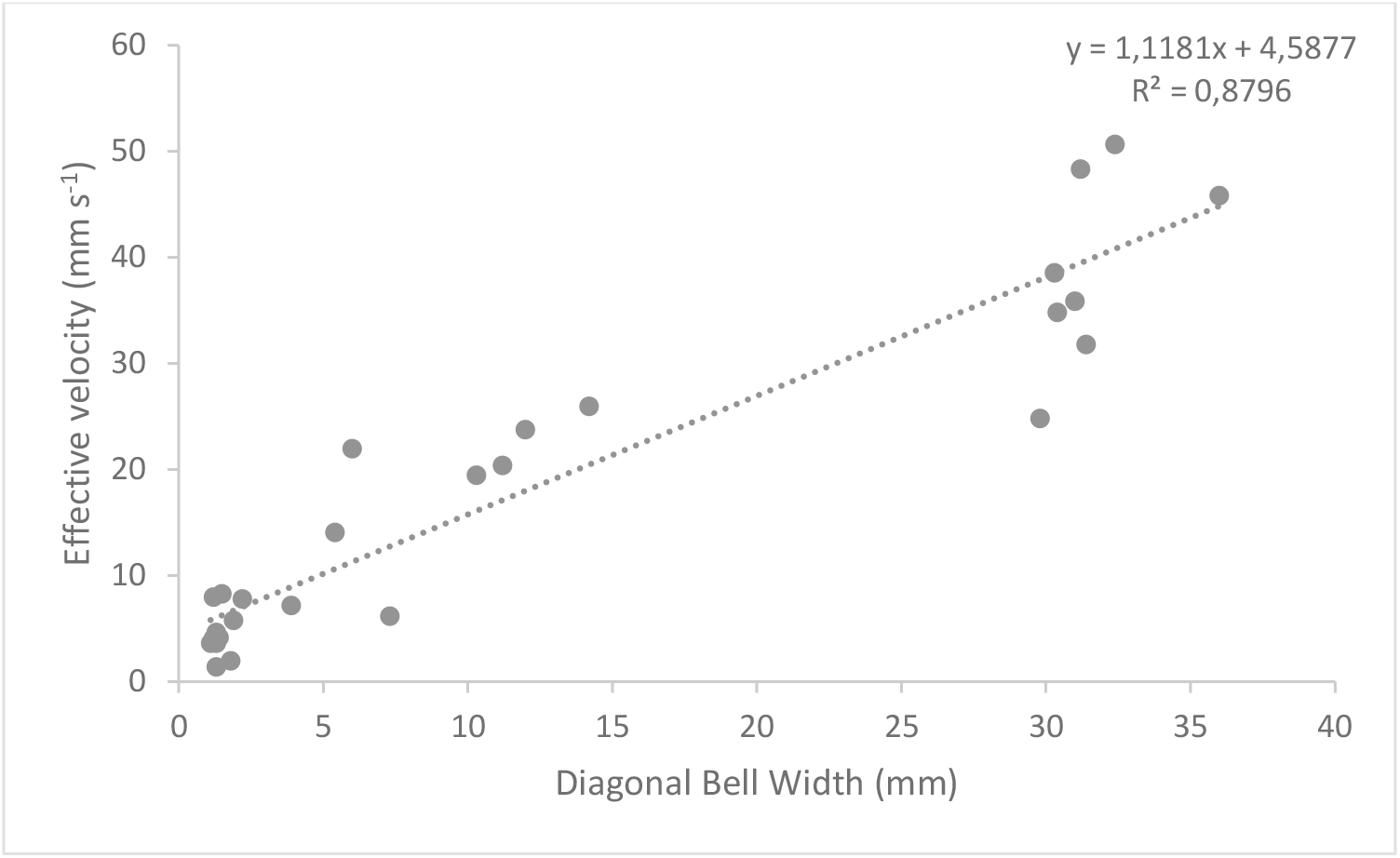
Effective velocity of *C. marsupialis* with increasing size (n=27).

### Proficiency

The calculated proficiency decreased following a potential function, with the highest values observed for the smallest juvenile stage (up to 9.47), and reaching a plateau in adults (1.37 ± 0.13, mean ± s.d., N=8)(Fig. 6).

**Fig. 6.**
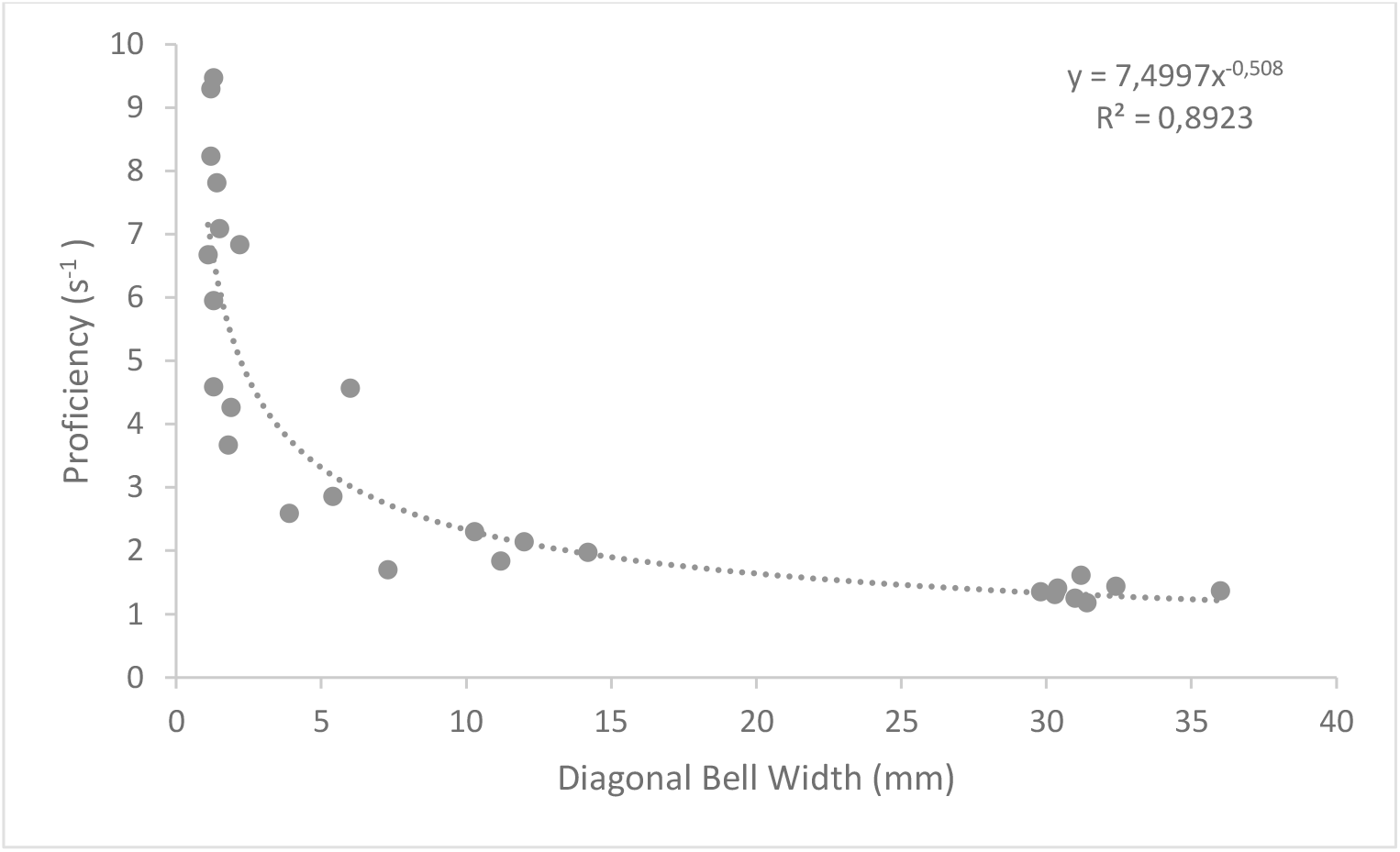
Swimming proficiency of *C. marsupialis* with increasing size (n=27).

### Surface current velocities and capability of *C. marsupialis* to overcome them

We analyzed a total of 850 short tracks of drifting buoys (X-Y m of track). The velocity of the currents in the study area varied depending on the month and the distance from the coast. At shoreline (0 m) the lowest values were obtained, ranging between 31.87 ± 3.82 mm s^-1^ in November and 58.01 ± 11.79 mm s^-1^ in May (mean ± s.e.m., N=32 and N=39, respectively). Grouping all months, the average current speed at this distance was 44.32 ± 3.05 mm s^-1^ (mean ± s.e.m., N=510). At 250 and 500 m, the values were higher and similar to each other, with mean values of 94.72 ± 10.82 mm s-1 and 107.71 ± 14.93 mm s-1, respectively, when considering all months. In both distances, the minimum values were obtained in November, while the maximum ones were recorded in July (41.18-42.47 mm s^-1^ and 120.09-159.24 mm s^-^1, at 250 and 500 m, respectively).

Taking into account these differences, we calculated the theoretical percentage of currents overcome by each stage, considering the months in which they are present. Specifically, we used May-July for the smaller specimens, July-August for the medium-sized specimens, and August-November for adults (see Fig. 7). We then compared the average speed of each specimen in each group with the calculated velocities of the buoys assigned to each period and distance The number of buoys used for calculating current speeds at 0, 250 and 500 meters, were: 250, 81 and 87 for the period May-July, 191, 48 and 60 for July-August and 260, 90 and 82 for August-November months, respectively.

**Fig. 7.**
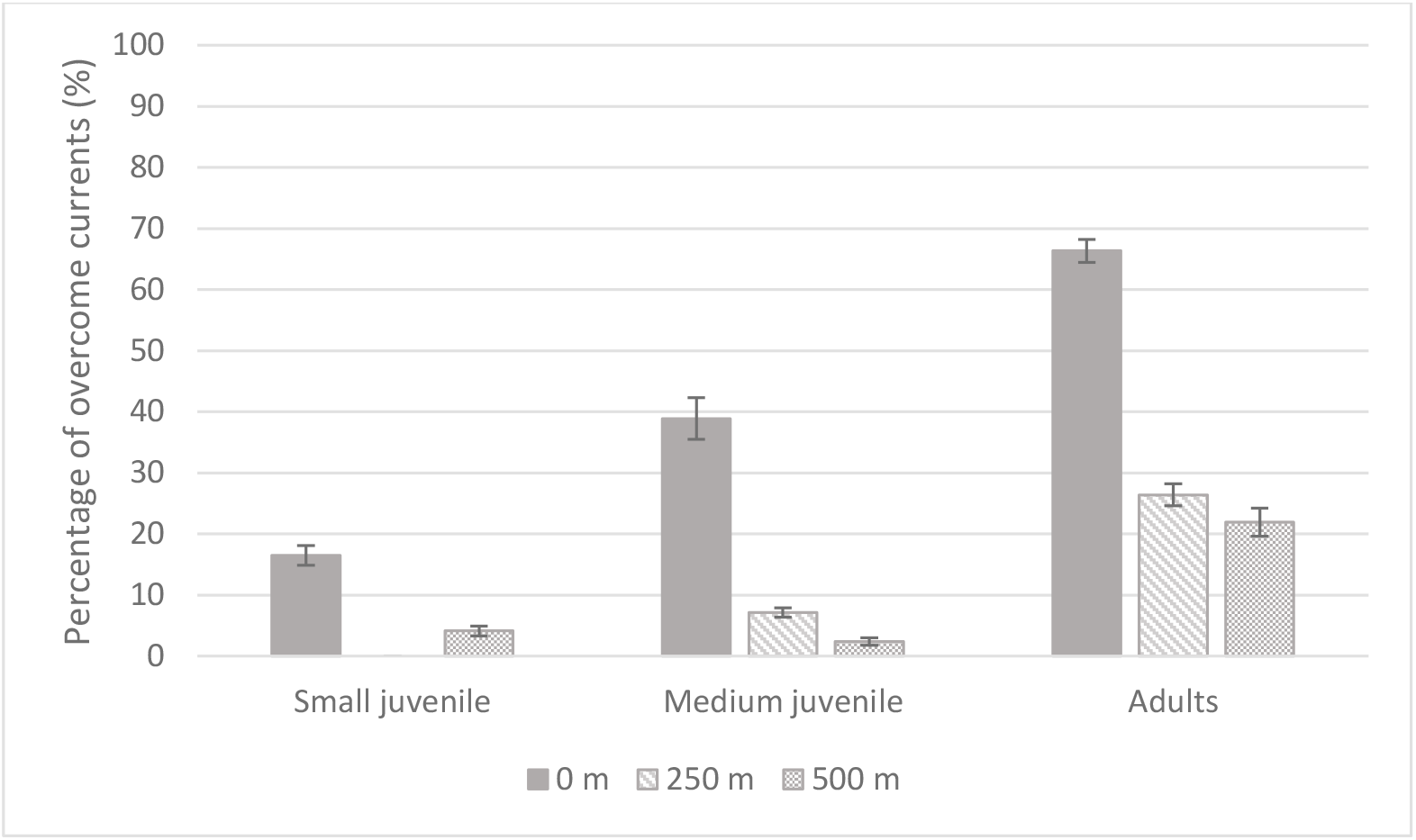
Percentage of current overcome of each size group of *C. marsupialis* in relation to their periods of presence and distance from the coast (mean ± s.e.m.)(n=12 for small juveniles, n=7 for medium juveniles and n=8 for adults).

While the adults are able to swim strongly enough to overcome almost 70% of currents at 0 m, this percentage decreases to below 27% at farther distances. For medium-size juvenile percentages are lower, not even reaching 40% on the shoreline. Small ones can practically not overcome currents at any distance (in fact its percentage is 0 at 250 m and the highest value, at 0 m, does not even reach 17%).

## DISCUSSION

### *Carbydea marsupialis* is a strong swimmer

We analyzed 27 specimens from 1.1 mm to 36 mm DBW, which showed an average swim speed of 22.74 ± 2.89 mm s^-1^ (mean ± s.e.m.), with higher speeds for larger individuals. This increase in swimming performance with size is a common trait among laboratory-tested cubozoans (Colin et al., 2013; Garm et al., 2007; Schlaefer et al., 2020; Shorten et al., 2005), but differs from the study of Schlaefer et al. on *C. fleckeri* in the field where the opposite relationship was found (Schlaefer et al., 2018). The authors suggested that this discrepancy may be due to the fact that the observed jellyfish in the wild were not swimming at their full capacity.

The average maximum speed recorded ranged from 18.64 mm s^-1^ for small juveniles (DBW ≤ 5mm) to 75.35 mm s^-1^ for adults (DBW ≥ 15 mm), with a speed of 40.98 mm s^-1^ for the medium-size class (5 ≤ DBW ≤ 15 mm). These velocities are comparable to those observed in other cubozoans of similar size range. For example, in trials performed in the laboratory, *T. cystophora* of 8-12 mm bell diameter (BD) and *C. bronzie* of 30-50 mm BD swam at maximum speeds of 3-4 cm s-1 and 7-8 cm s-1 against a 1-1.5 cm s-1 current, respectively (Garm et al., 2007). Meanwhile *C. sivickisi* with IPD (inter pedial diameter, a distance between opposite pedalia at the level of the bell turn-over, a measure slightly larger than DBW, Acevedo et al., 2019) ranging from 4 to 11 mm 4.9 cm against currents of 3-18 cm s^-1^ (Schlaefer et al., 2020). For *C. fleckeri*, Schlaefer et al. 2018 reported a maximum speed of 6.5 cm s^-1^ for a specimen of 4 cm IPD and Hamner (1995) described speeds about 8 cm s^-1^ for jellyfish of 30-100 mm BD.

The calculated EDI per size groups varied from 0.51 in small specimens to 0.90 in adults. Although based on speeds and velocities rather than distances and displacements, our EDI would be comparable to the NGDR (net to gross displacement ratio, i.e. displacement/distance)(Buskey et al., 1993; Vidal et al., 2018). In Buskey et al., the crustacean *Artemia salina* showed a variation in its NGDR from 0.11 to 0.55 with increasing nauplius stage, and initial nauplius stages of different copepod species presented low NGDR values (0.12-0.35) that increased up to 0.65 for second nauplius stage copepods. On the contrary, Vidal et al. (2018) registered decreasing NGDR values for the paralarvae of the squid *Doryteuthis opalescens* (from 0.63 to 0.36 for individuals of 2.65 to 9.81 mm mantle lengths, respectively). However, in their study, they applied a current of 1 cm s^-1^ that smaller individuals were not able to overcome, so they showed relatively long horizontal displacements due to the drift. The bigger squids, on the other hand, spent most of the time hovering (when doing so in the wild they remain in the same area, e.g. areas of high food availability), which makes the relative path displacement considerably reduced. To the best of our knowledge, there are no studies that determine the NGDR of cubozoan species.

The observed proficiency trend in *C. marsupialis* according to size is also consistent with those obtained in other box jellyfish such as *C. fleckeri* and *C. bronzie* (Colin et al., 2013), as well as the hydromedusa *Sarsia tubulosa* (Katija et al., 2015). As the bell diameter increased, the swimming proficiency decreased. Proficiency is related to foraging strategies. Species that forage as cruising predators have associated a lower swimming proficiency whereas ambush foraging species swim more proficiently (Dabiri et al., 2010). Adults of *C. marsupialis* showed a proficiency value of 1.37 ± 0.13 (mean ± s.d., N=8), which would be consistent with a cruising foraging strategy (Colin et al., 2013; Kiørboe, 2011) as previously stated (Acevedo et al., 2013).

Regarding the surface currents present in the study area and a comparison with the average speeds calculated for the different size groups, only adults could –theoretically– overcome them in a remarkable way. This result could be key to understanding their spatiotemporal distribution. On the coast of Dénia (Western Mediterranean) juveniles and adults are usually found in different areas some kilometers apart. Whereas adults are present in high productivity areas, juveniles have a more dispersed distribution (Bordehore et al., 2020a; Canepa et al., 2017), far from these highly productive areas.

These results support the hypothesis that adults have strong enough swimming abilities to select their habitat, whereas juveniles rely on drifting currents and thus reflect the advection pattern caused by these currents. Nevertheless, it would be necessary to conduct experiments using currents to verify it completely.

The methodology developed in this paper allows the acquisition of high-quality swimming speed measurements for specimens of size ≥ 1 mm with conventional video recordings. In addition, it allows the animal to swim freely and, therefore, represents a good approximation of its behavior in the wild. This information is increasingly valuable because including empirically derived jellyfish behavior in particle tracking models that are used to forecast the timing and magnitude of jellyfish proliferation near major tourist areas, aquaculture facilities or power plants, makes them more realistic (Fossette et al., 2016). It should be noted, that the last months of the bathing season in the Mediterranean (August-September) overlaps with the months when *C. marsupialis* reaches its adult stage (Boero and Minelli, 1986; Bordehore et al., 2020a; Canepa et al., 2017) so knowing its presence in advance can be very useful. Although its sting can be considered of moderate intensity (Kokelj et al., 1992; Peca et al., 1997) with cutaneous affection mainly, it can further generate systemic effects in an unknown percentage of the population (Bordehore et al., 2015b).

## Acknowledgements

We are grateful for the collaboration of Fundació Baleària, Marina El Portet de Dénia and Marina de Dénia. We also thank the students and volunteers from UA and the Marine Laboratory UA-Dénia for their support on jellyfish and drifting buoys samplings: Lucía Portilla, Carlos de Juan, Cristina Alonso, Lara Sánchez, Neus Figueras, Vicente Bernabeu, Roberto Cabria, Patricia Rigio, Luis Arechavaleta, M^a^ José Vargas, Sofía Capellán, Laura Avivar, Beatriz Rubio, Juan Pablo Izaguirre, Ainara Ballesteras and Héctor Gutiérrez, as well as, JA. Yañez-Dobson for creating the figure 1.

